# Multi-Methodological Characterization of Sleep Deprivation: From Standard EEG Power Spectra to Aperiodic Dynamics in Humans and Mice

**DOI:** 10.64898/2026.09.08.750125

**Authors:** Thomas Kroker, Lena Puder, Omid Abbasi, Sara Ghiasi, Clemens Krug, Ida Wessing, Mohammad Ali Salehinejad, Tillmann Ruland, Judith Alferink, Udo Dannlowski, Philipp Ritter, Joachim Gross

## Abstract

Sleep deprivation is a potent, rapid-acting therapeutic intervention for major depressive disorder, yet its underlying neural mechanisms remain poorly understood, hindering the development of predictive biomarkers. Here, we systematically characterize the electrophysiological signatures of prolonged wakefulness using a multi-methodological approach across three independent datasets in humans and mice. By integrating standard power spectral density analysis with aperiodic component fitting (SpecParam) and highly comparative time-series analysis (HCTSA), we identified robust cross-species biomarkers of sleep pressure. Machine learning models revealed that theta power is the most consistent feature for differentiating control and sleep deprivation states, achieving up to 90% classification accuracy. Sleep deprivation significantly increased the spectral offset - suggesting global cortical hyperexcitation - while simultaneously steepening the spectral slope. We interpret this simultaneous shift as a state uncoordinated state of hyperexcited and inefficient neural processing. These findings establish reproducible EEG markers of sleep deprivation that transcend species. Given the clinical utility of wake therapy, we propose that prefrontal theta power and spectral offset/slope may serve as mechanism-based predictors of therapeutic response. Our results provide a framework for the clinical validation of these biomarkers, potentially enabling personalized chronotherapeutic interventions for psychiatric disorders.

## Introduction

Sleep deprivation has now become a well-accepted treatment option for depression due to its rapid mood-improving effects (1,2), however the underlying mechanisms remain poorly understood. The identification of biomarkers that can predict which patients will respond and which will not is an important step in that direction. The most interesting candidate is the MEG/EEG power in the theta frequency band (4-8 Hz) which has been proposed before as a potential biomarker for treatment response to wake therapy (3), but the evidence is conflicting (4,5). Sleep pressure and theta power play an important role in the effectiveness of wake therapy, however some studies suggest that excessive theta responses to wake therapy predict poor treatment response (6), while other studies suggest the opposite (3). Importantly, theta activity is associated with sleep pressure (7,8), that builds up during the day and further increases after sleep deprivation. With consideration for the application of wake therapy in depressed patients (9,10), it is crucial that sleep deprivation is related to neural plasticity and synaptic strength (11). In this regard, sleep deprivation can induce an enhanced global synaptic strength indicated by higher wake EEG theta activity. However, only responders show an association with neuroplasticity (i.e., increased theta power) whereas non-responders do not (4,5). A further disorder, for which it is appealing to understand sleep deprivation better is non-organic insomnia. This is because the evidence regarding the associations between the inability to sleep, sleep pressure and hyperarousal with theta is also contradictory. Some studies suggest an increase in theta activity in patients with insomnia linking this to hyperarousal during wakefulness (12,13), whereas other studies find the opposite pattern (14).

Beyond theta power, further measures worth considering as potential biomarkers of sleep deprivation are the offset and slope of the MEG/EEG power spectrum (15). This approach capitalizes on the fact that the MEG/EEG power spectrum consists of periodic and aperiodic components. The power spectrum was decomposed into its periodic and aperiodic (offset and slope) components. The offset of the power spectrum has been linked to general neural activity and a higher offset indicates stronger neural activity (16). The slope of the power spectrum has been associated with the balance of excitation and inhibition (E/I balance; (17)) with steeper slopes (dominance of low frequencies) indicating relatively increased inhibition and flatter slope with increased excitation (18,19). Furthermore, the slope of the power spectrum has also been employed as a measure for neural plasticity (11). Offset and slope are also sensitive to sleep deprivation (20). In particular, a recent study showed an increased offset following sleep deprivation (21). This increase in cortical responsiveness (i.e., elevated offset) was associated with reduced subjective alertness. The authors interpret this increase in neural activity as being due to continuous upregulation of neural firing during prolonged wakefulness.

To improve our understanding of the neural mechanisms involved in sleep deprivation, we performed analyses on discovery, replication and combined EEG datasets. To this end, we analyzed two human resting-state EEG datasets. The first (21) was measured by a Chinese group at the Southwest University in Chongqing (discovery dataset) and the second (replication dataset) was gathered in Germany at the Leibniz Research Centre for Working Environment and Human Factors in Dortmund (22). In a second step, we calculated our analyses over both datasets, to extract the most robust biomarkers of sleep deprivation in human resting-state EEG. To make our analysis even more robust and show its neurobiological consistency and plausibility across species, we performed a convergent analysis with the same methods on mouse data provided by a Korean group (23). This approach seems especially tempting since a lot of the above described mechanisms have been discovered in the mouse model also by making use of invasive measurements, i.e., synaptic plasticity induced by sleep deprivation (24–26) or upregulated neural activity after sleep deprivation (27,28). It is well known that sleep deprivation cannot only induce mania in humans, but also in mice. In this regard, particularly prefrontal changes in synaptic plasticity via dopaminergic pathways following sleep deprivation seem to be responsible for inducing mania-like behavior in mice. Furthermore, inflammatory and microglial processes appear to be involved in these behavioral changes, i.e., increased social activity, exploration time and total distance travelled(25,26).

For our analysis we combine a comprehensive set of features with univariate and multivariate statistical analysis. Our set of features consisted of standard EEG spectral power, EEG spectral power corrected for aperiodic activity, estimates of aperiodic activity (offset, slope) and about 7000 additional features from the highly comparative time-series analysis (HCTSA). The HCTSA features characterize the EEG time series across many domains including its correlation, complexity, distribution etc.(29). For univariate analysis we employed channel-wise T-statistics. For multivariate analysis we used cross-validated support-vector machines.

We hypothesize that theta power will increase after sleep deprivation compared to the sleep control condition as it is well known that theta activity increases with sleep pressure (8,30). This increase will be reflected in univariate statistical analyses, as indicated by strong effects in the theta range. Furthermore, multivariate machine learning analyses will precisely differentiate between sleep deprivation and the control condition in the theta band, as indicated by high classification accuracies in this frequency band. Regarding the slope and offset, we expect to replicate previous findings showing an increase in the offset of the power spectrum(11,21) as well as an increase in the spectral slope with sleep deprivation (11). Due to the exploratory nature of the highly comparative time-series analysis (HCTSA), we do not formulate any hypotheses.

## Results

To identify robust EEG biomarkers of sleep deprivation, we analyzed resting-state wake EEG data from a discovery dataset (n=65) and a replication dataset (n=31), as well as a combined dataset (n=96). By utilizing only the eyes-open data, we ensured that our analysis was not confounded by periods of microsleep. Our analysis followed a pre-defined pipeline with a multi-methodological approach (Figure 1) to systematically test markers, moving from standard spectral measures to advanced aperiodic and time-series-based features. For the univariate tests, we used the entire dataset (i.e., the Chinese and German datasets combined). In the multivariate machine learning approach, however, we trained the model using only the Chinese data and tested it using the German and combined datasets. For the analysis of the mouse data (n=9), we used the same analysis methods with some adjustments to the specificities of the particular dataset.

**Figure 1.**
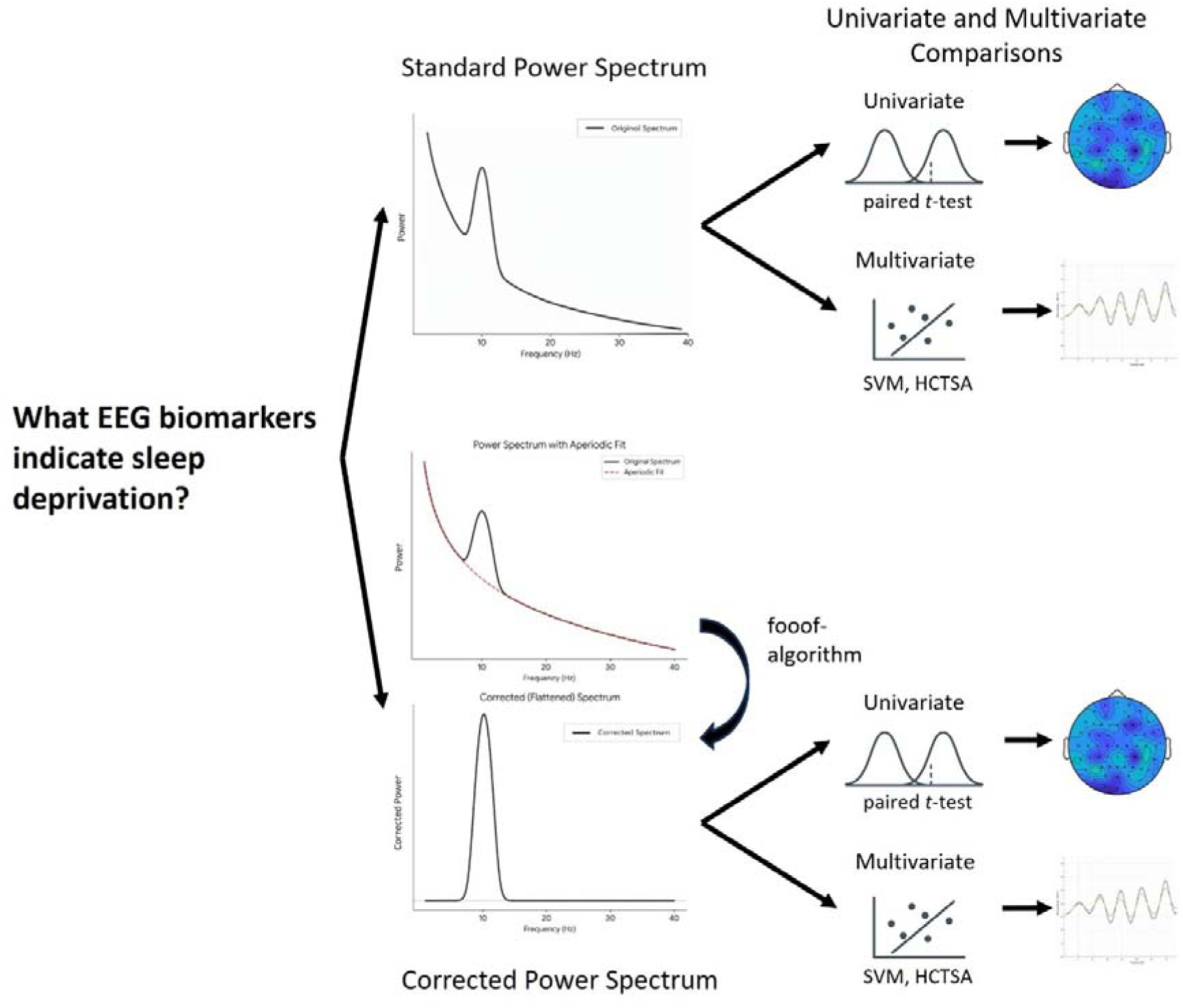
Schematic illustration of the applied methods and analyses. This includes the standard and corrected power spectra, which were analyzed with univariate as well as multivariate methods (support vector machine).

### Features of the standard power spectrum

First, we calculated univariate statistics for the standard power spectrum from 0-100 Hz over both datasets to differentiate between sleep deprivation and normal sleep. The cluster-corrected t-tests comparing both sleep conditions revealed a maximal t-cluster-statistic of - 126.89 at 6.25 Hz (cluster *p*-value < 0.001) indicating the strongest effect across all frequencies in the theta band. Theta power is therefore significantly stronger after sleep deprivation compared to normal sleep. Other significant effects were found at about 3.25 Hz and between 20-30 Hz. To assess discriminability, we trained a linear support vector machine (SVM) classifier on single-frequency bins using 10-fold cross-validation (Figure 2). We trained three classifier. First, we trained a classifier on the discovery data and assessed performance (classification accuracy and AUC (area under curve) on the test folds of the 10-fold cross-validation. Second, we trained a classifier on the discovery data and assessed performance on the replication data that was recorded on a different site with a different EEG system. Third, we trained a classifier on the combined data and, again, assessed performance on the test folds of the 10-fold cross-validation. Classification accuracy peaked in the theta range, 90.14% (*AUC* = 0.95) at 6.5 Hz in the discovery data, 90.00% (*AUC* = 0.95) at 4 Hz in the replication data and 84.15% (*AUC* = 0.90) at 6.25 Hz in the combined dataset (Figure 2C). Thus, theta oscillations reliably differentiated sleep deprivation from normal sleep across datasets.

**Figure 2.**
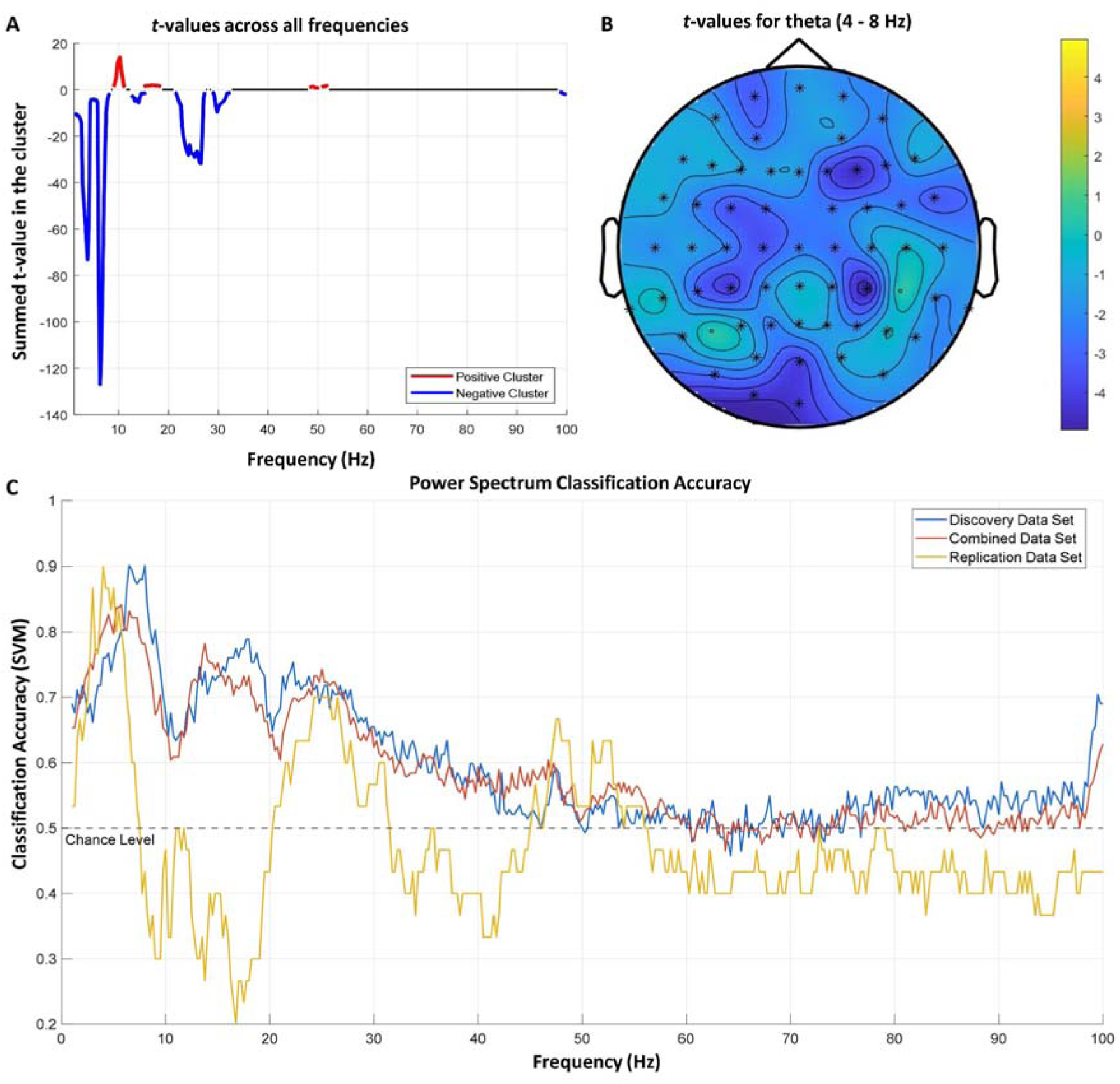
A. t-values for positive and negative clusters in each frequency differentiating between normal sleep and sleep deprivation. B. Topography plot indicating t-values testing normal sleep against sleep deprivation in the theta frequency range. C. Accuracy with which sleep deprivation can be differentiated from normal sleep across all frequencies. A linear SVM (support vector machine) model with 10-fold cross-validation was used separately for each frequency bin. Accuracy is shown for three models. First, accuracy for the discovery data (blue line), second, accuracy of the discovery model tested on the replication data (yellow line) and, third, accuracy on both data sets combined (red line).

### Separating periodic and aperiodic activity

Next, we removed the aperiodic component from the power spectrum to investigate, to what extent the remaining periodic components of the power spectrum change differ after normal sleep and sleep deprivation. The results in the corrected power spectrum are similar to the ones in the normal power spectrum (maximal *t*-cluster-statistic =-143.70 at 6.25 Hz; cluster *p*-value < 0.001; Figure 3A-B). Importantly, this result demonstrates that sleep deprivation has a significant effect on periodic theta power irrespective of aperiodic activity. However, some additional effects can be seen in the corrected power spectrum. At the statistical level, a pronounced difference is the positive cluster in the gamma range (peaking at 53 Hz, *p*-value < 0.001; *t*-cluster-statistic = 70.42), indicating higher gamma power after normal sleep compared to sleep deprivation. However, this effect is not present in the multivariate analysis, where the classification accuracy of this frequency is not superior compared to neighboring frequencies. Again, the most precise classification can be reached when the theta frequency is used (Figure 3C), although not as strong as in the uncorrected spectrum. Classification accuracies reached their maximums of 77.46% (*AUC* = 0.87) at 6.25 Hz in the discovery data, 90.00% (*AUC* = 0.95) at 5.75 Hz in the replication data and 76.23% (*AUC* = 0.87) at 26.25 Hz in the combined dataset (peak in theta range at 6.5 Hz with 74.91% accuracy and AUC = 0.84).

**Figure 3.**
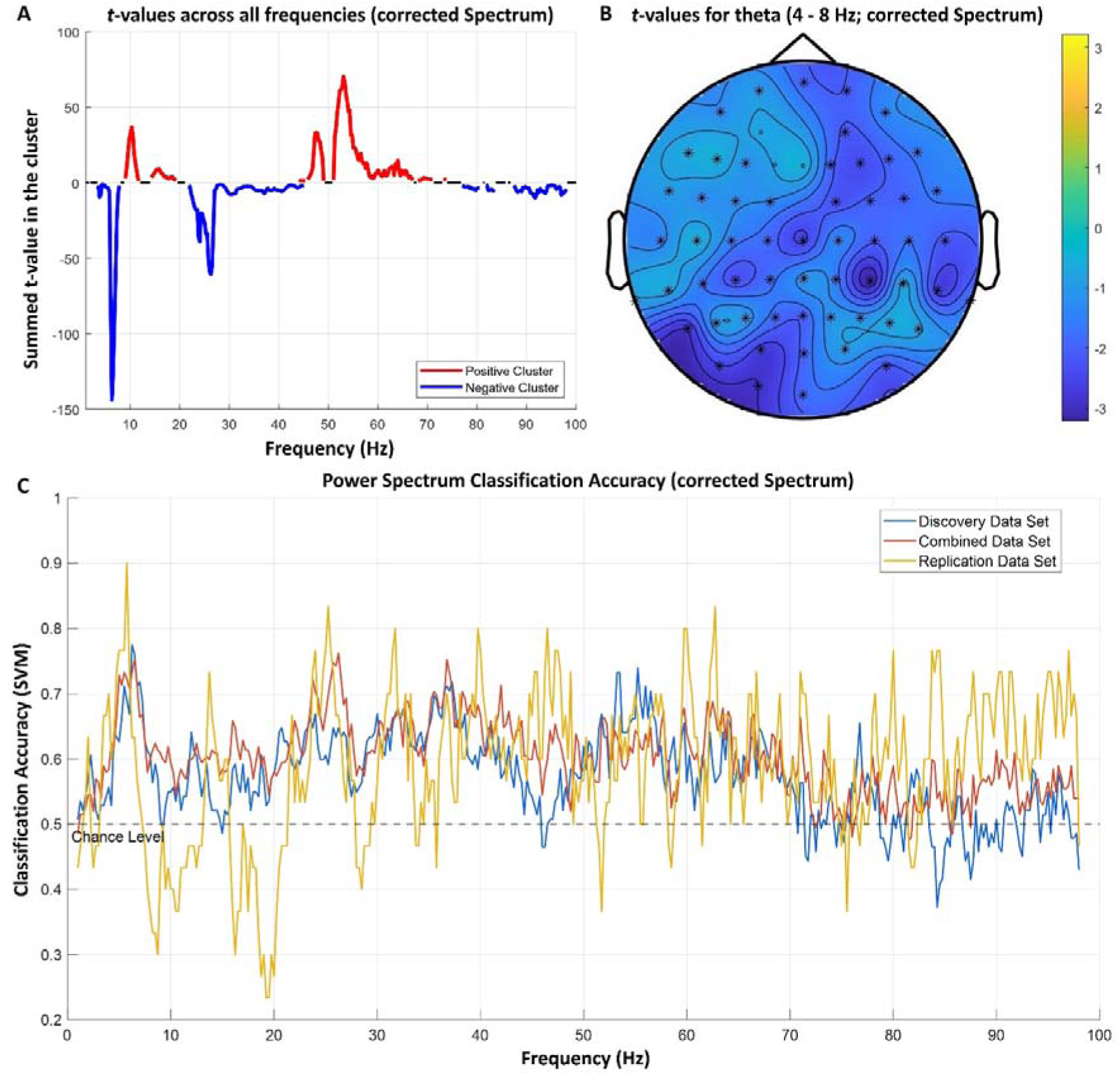
A. t-values for positive and negative clusters in each frequency differentiating between normal sleep and sleep deprivation in the power spectrum after removing the aperiodic component. B. Topography plot indicating *t*-values testing normal sleep against sleep deprivation in the theta frequency range (corrected power spectrum). C. Accuracy with which sleep deprivation can be differentiated from normal sleep across all frequencies in the corrected power spectrum. A linear SVM (support vector machine) model with 10-fold cross-validation was used separately for each frequency bin. Accuracy is shown for three models. First, accuracy for the discovery data (blue line), second, accuracy of the discovery model tested on the replication data (yellow line) and, third, accuracy on both data sets combined (red line).

**Figure 4.**
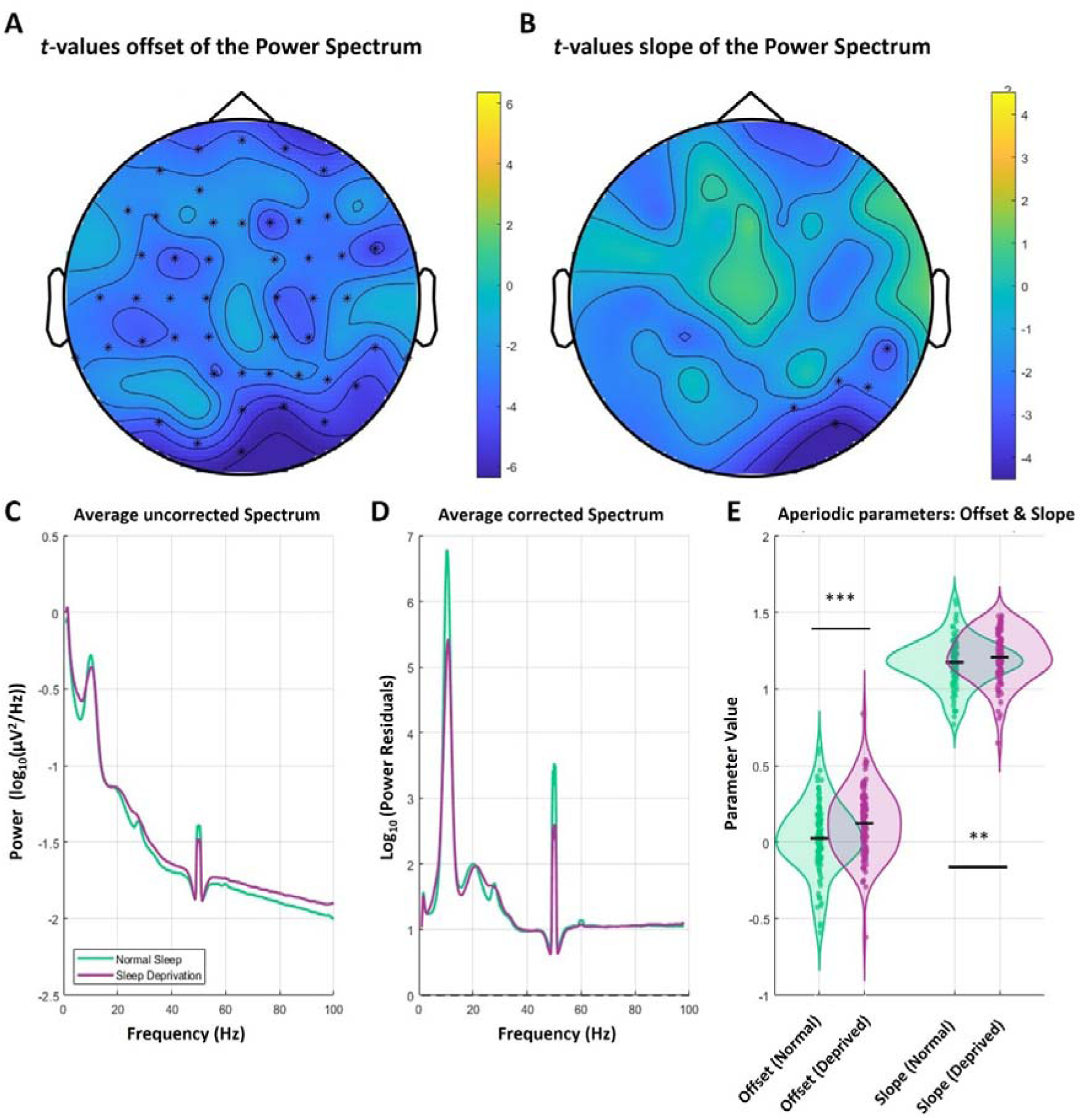
A. Topography plot indicating t-values testing offset of the corrected power spectrum after normal sleep versus sleep deprivation. B. Topography plot indicating t-values testing slope of the corrected power spectrum after normal sleep versus sleep deprivation. C. Uncorrected power spectrum after sleep deprivation and normal sleep. D. Corrected power spectrum after sleep deprivation and normal sleep. E. Offset and slope of the corrected power spectrum after normal sleep and sleep deprivation.

Offset and Slope

While the previous analysis focused on the periodic component of the power spectrum, we next investigated the aperiodic component. Comparing the offset and slope of the power spectrum revealed strong effects, particularly for the offset, which was present across electrodes spread across the entire scalp. However, the strongest effects of sleep deprivation are over the posterior electrodes as indicated by the *t*-values. Regarding the slope (exponent), effects between the sleep conditions only occurred over occipital electrodes. To test global effects on slope and offset, we averaged these overall electrodes and used a paired t-test to compare the sleep conditions, which revealed effects on the offset (p-value < 0.001; t-cluster-statistic =-57.88) and on the slope (*p*-value < 0.001; *t*-cluster-statistic =-9.21). Offsets were higher after sleep deprivation, whereas slope increases were predominantly occipital and may vary with fitting choices. The increased offset suggests higher general neural activity after sleep deprivation. Importantly, the SpecParam-algorithm returns the exponent of the power spectrum, i.e., the higher exponent indicates a steeper spectral slope after sleep deprivation. This signals a shift towards slower frequencies suggesting relatively enhanced inhibition in the E/I balance. Classification accuracies for offset were 66.90% (*AUC* = 0.75) in the discovery data, 70.30% (*AUC* = 0.76) in the replication data and 70.00% (*AUC* = 0.67) in the combined dataset. Slope classification accuracies reached 48.59% (*AUC* = 0.53) in the discovery data, 60.00% (*AUC* = 0.60) in the replication data and 53.47% (*AUC* = 0.56) in the combined dataset.

### HCTSA

We extended our previous analysis to a comprehensive set of time-series features to investigate if we can find a specific characteristic of sleep-deprivation induced changes in the resting-state EEG with a stronger effect size than theta power. Interestingly, none of the more than 7000 features included in the HCTSA toolbox outperformed theta power. The best discriminating feature between sleep deprivation and normal sleep was “CO_Embed2_Basic_tau_incircle_2”, which can be described as a measure of variance or complexity of the signal (Figure 5). The positive *t*-values on the topographies indicate a reduced variance after sleep deprivation compared to normal sleep (*p*-cluster < 0.001; *t*-cluster-statistic = 63.56). The SVM-accuracy differentiating between the sleep conditions reached 78.16% (*AUC* = 0.80) for the discovery set, 73.33% (*AUC* = 0.82) for the replication data set and 75.74% (*AUC* = 0.80) for the combined dataset.

**Figure 5.**
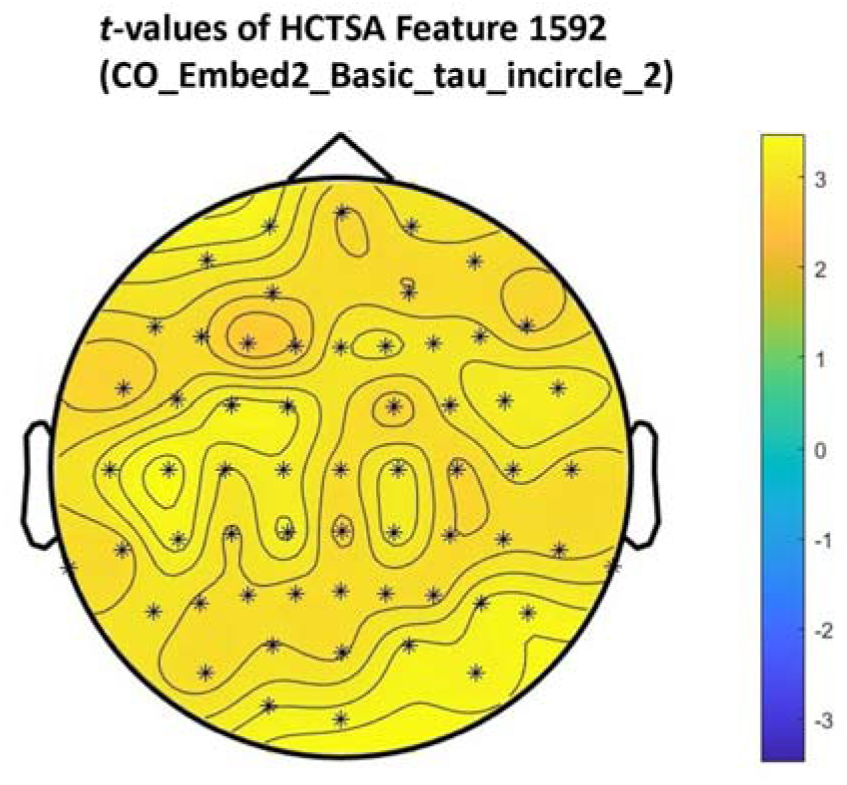
Topography plot indicating *t*-values testing HCTSA feature 1592 (CO_Embed2_Basic_tau_incircle_2) after normal sleep versus sleep deprivation.

In summary, our analysis reveals that theta power is the most reliable proxy for the neuronal effects of sleep deprivation. Significant effects are also evident (albeit with a lower effect size) in the offset and slope of the aperiodic component of the power spectrum.

### Complementary analysis of mouse data

Finally, we applied our univariate and multivariate analysis approach to resting-state EEG data recorded in mice following sleep deprivation and normal sleep. We aimed to investigate to what extent our previous results generalize across species.

Analogous to the human data, we tested the effects of sleep deprivation in the normal and corrected power spectrum. The cluster-corrected t-tests again showed strong effects between the two sleep conditions (maximal t-cluster-statistic =-57.94 at 8.5 Hz; cluster p-value < 0.001; Figure 6A). In contrast to the human data, the machine learning approach was applied within each mouse, as only nine mice were measured, but with a considerably longer duration of two hours. Accordingly, a lot more data per mouse was available compared to the human EEG data. Therefore, we used SVM-models for the classification of individual data segments to the two conditions (normal sleep or sleep deprivation), In this approach, classification accuracy peaked in the theta range as well: 89.00% at 7 Hz (AUC = 0.95; Figure 6C).

**Figure 6.**
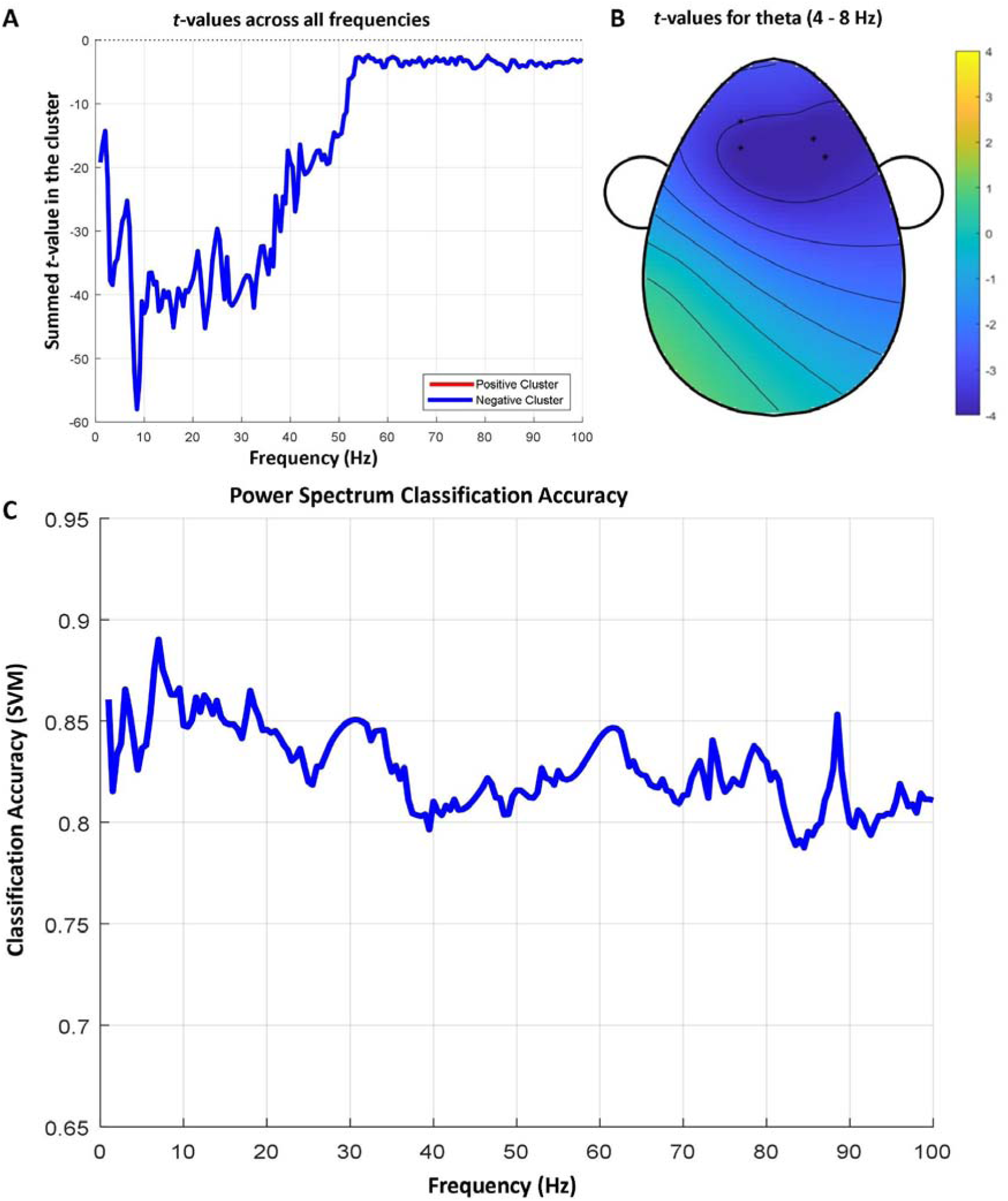
A. t-values in each frequency of the standard power spectrum differentiating between normal sleep and sleep deprivation in mice. B. Topography plot indicating t-values testing normal sleep against sleep deprivation in the theta frequency range for the mouse data. C. Accuracy with which sleep deprivation can be differentiated from normal sleep across all frequencies. A linear SVM (support vector machine) model with 10-fold cross-validation was used separately for each frequency bin.

In the corrected power spectrum, the strongest statistical effect was found at 52.50 Hz where the cluster-mass reached 103.74 (Figure 7A). The heaviest negative cluster was found at 9 Hz with a cluster-mass of-76.14 (*p* < 0.001). Peak classification accuracy of the SVM was reached again at 7Hz with 91.45% (*AUC* = 0.96; Figure 7C).

**Figure 7.**
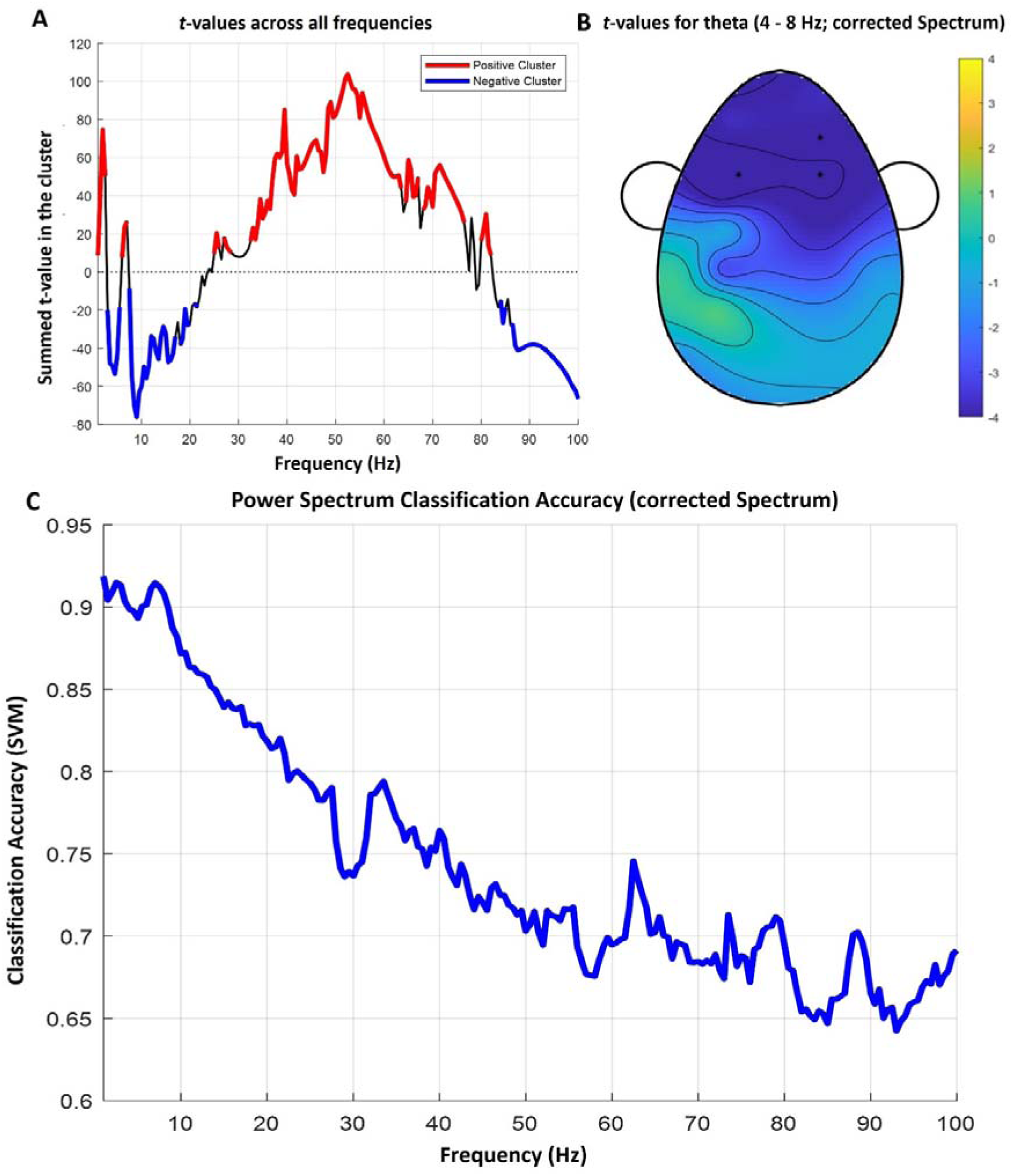
A. t-values in each frequency of the corrected power spectrum differentiating between normal sleep and sleep deprivation in mice. B. Topography plot indicating t-values testing normal sleep against sleep deprivation in the theta frequency range (corrected spectrum) for the mouse data. C. Accuracy with which sleep deprivation can be differentiated from normal sleep across all frequencies. A linear SVM (support vector machine) model with 10-fold cross-validation was used separately for each frequency bin (corrected spectrum). In contrast to the human data, this approach was calculated within each mouse, as only nine mice were measured, but with a considerably longer duration of two hours. Accordingly, a lot more data per mouse was available compared to the human EEG data.

**Figure 8.**
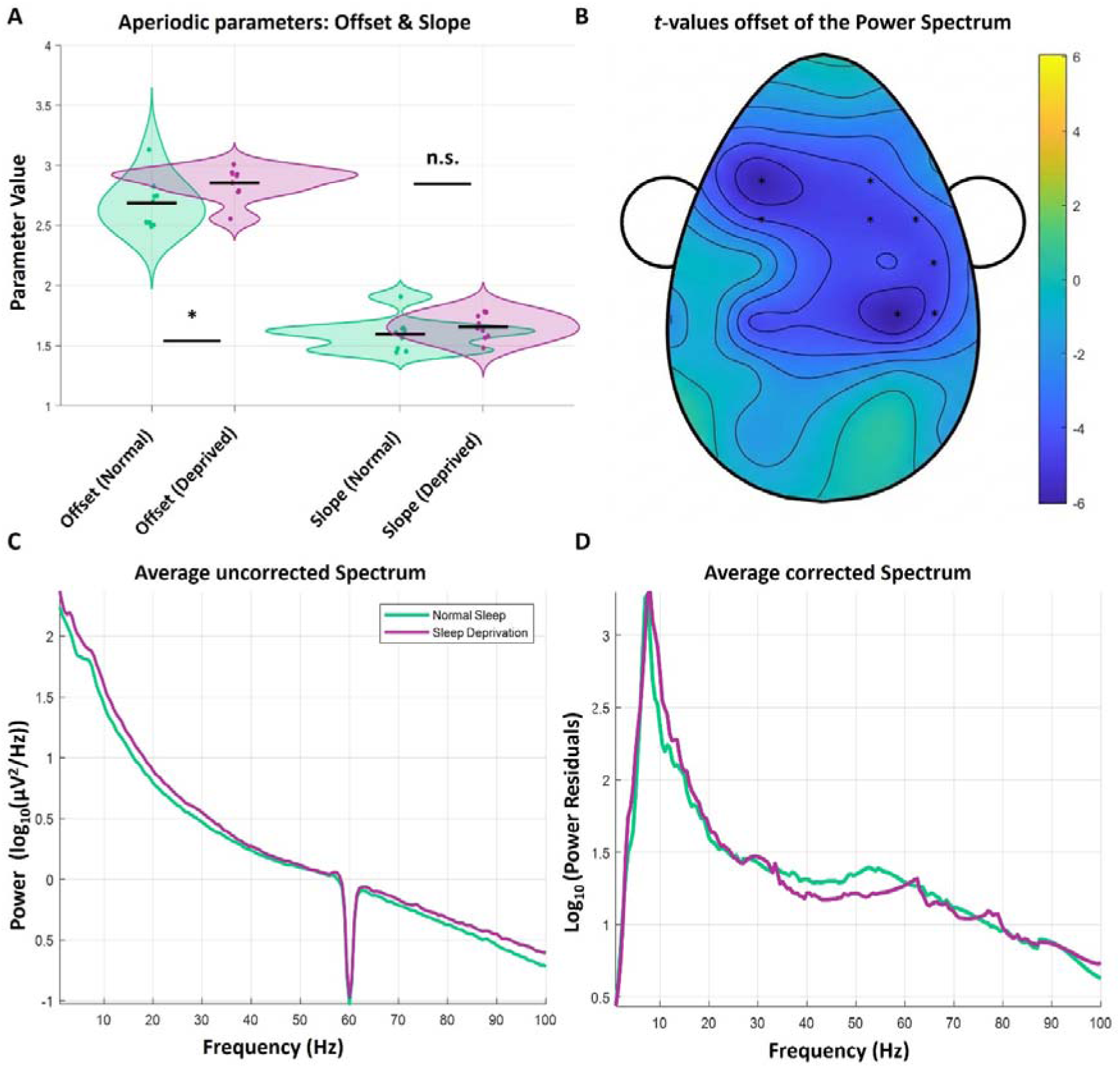
A. Offset and slope of the power spectrum after normal sleep and sleep deprivation. B. Topography plot indicating t-values testing offset of the corrected power spectrum after normal sleep versus sleep deprivation in mice. C. Uncorrected power spectrum after sleep deprivation and normal sleep. D. Corrected power spectrum after sleep deprivation and normal sleep.

Offset and Slope

Differences were found in offset (*t*(8) =-2.76, *p* = 0.025) but not for slope (*t*(8) =-0.02, *p* = 0.985) Cluster-based statistics revealed a significant cluster for offset (*p*-cluster = 0.019; *t*-cluster-statistic =-33.45), but not for slope (*p*-cluster = 0.109, *t*-cluster-statistic =-3.05) in the mouse data. In the machine learning approach accuracies of 66.67% (AUC = 0.69) were reached for offset and 44.44% (AUC = 0.48) for slope.

## Discussion

### Standard power spectrum

We found a strong association between theta power and sleep deprivation, which is well in line with previous literature showing an increase in theta power with prolonged wakefulness (22,30,31). This increase in theta power was associated with a decrease in cognitive performance, such as attention, working memory or reaction times. The effects were strongest over parietal and occipital electrodes, however they are present across the whole scalp. The novelty and strength of our study lies in our ability to robustly differentiate between sleep deprivation and normal sleep across different types of univariate and multivariate analyses and biological species using theta power. One of the main goals of this study was to take a step back from wake therapy for depression in order to understand how activity patterns in the brain change after prolonged wakefulness, and how these changes could improve depressive symptoms. Frontal theta activity is particularly altered in affective disorders, tending to be reduced in depression (32) and increased in mania (32–34). However, results are inconclusive regarding depression, some studies suggest increases in theta power, while others show a decrease (35). Furthermore, the same heterogeneity applies to treatment responses (36), but results tend to show increased (prefrontal) theta power after non-medication treatment response (37–39), while the pattern is the opposite for treatment response to medication (40,41). This strongly suggests that different mechanisms of action work for different patients and potential subtypes of depression. This is not surprising given the heterogeneity of depressive trajectories across patients, and suggests that more studies focusing on predicting individualized treatment responses using biomarkers are needed. Combining these previous results with our findings could help here. One might conclude that reduced (prefrontal) theta power is a potential biomarker for the response to wake therapy, since our analysis demonstrates increased theta power following prolonged wakefulness. Therefore, depressive patients with reduced prefrontal theta power may be more likely to respond to wake therapy, as sleep deprivation could restore prefrontal neural plasticity accompanied by a boost in theta power (5). As our findings in healthy subjects are not generalizable to clinical populations, future studies should validate these proposed markers in clinical cohorts to advance personalized interventions.

### Periodic and aperiodic activity

After correcting the power spectrum for aperiodic activity, theta power was still the most reliable frequency band in differentiating between the two sleep conditions as indicated by our univariate and multivariate analyses. Importantly, however, the corrected spectrum showed effects in the beta and gamma band particularly, which were less evident in the standard power spectrum. The evidence regarding changes in beta and gamma power is sparse and partially contradictory, making the interpretation of our results difficult. Studies tend to show a decrease in beta and gamma (42–44) after sleep deprivation, while other studies suggest an increase in beta (45). The decrease in gamma, however, seems to be more stable and we also found this reduction in gamma power after sleep deprivation. Regarding beta we found an increased power after sleep deprivation, which rather contradicts the existing literature. Nevertheless, it is important to note that only one of the above-mentioned studies corrected for aperiodic activity, so that it could be that the effects we found in beta and gamma can hardly be detected without this correction. Conversely, it is conceivable that the correction for aperiodic activity gives rise to a particular artifact in higher frequency ranges. Because if the statistical effects in the gamma band provide actual information these effects should generalize to the machine learning approach, which is barely to not at all the case. Accordingly, it is strongly discussed which frequency range should be fitted to identify periodic and aperiodic components properly (15,46,47). Nevertheless, theta power remains the most reliable frequency band across the corrected and uncorrected power spectra to differentiate between the sleep conditions.

### Offset and Slope

Additionally, the SpecParam-algorithm returns the offset and the slope of the power spectrum. The offset of the power spectrum is thought to reflect general neural activity, i.e., how many neurons fire simultaneously (16). We found the offset to be increased after sleep deprivation, which is clearly in line with the findings of other groups (21,48) and indicates a global cortical hyperexcitation following prolonged wakefulness. The neurobiological basis of this result can be interpreted as hyperexcitation of the brain induced by prolonged wakefulness, which could potentially induce mania in individuals at risk (49,50). The increased exponent (i.e., steeper slope) after sleep deprivation compared to normal sleep is more complicated to interpret, as this steeper slope means a shift towards slow oscillations. Conversely, the proportion of high-frequency oscillations in the power spectrum decreases relative to slow waves. At first glance, this seems puzzling, as an enhanced offset is usually taken to mean increased overall neural activity in the brain. However, a steeper slope with more slow-wave activity is usually taken to mean increased inhibition compared to excitation (18,19). A potential explanation for finding both at the same time could be that increased offset and slope simultaneously could reflect a state of uncoordinated neural overexcitement, just like an overheating engine, that is running fast but inefficient. Because this condition may reflect an abnormal E/I balance, indicating increased neural noise and inefficient information processing (51). Furthermore, this pattern with a relatively increased offset and slope at the same time is associated with certain neuro-developmental and neuropsychiatric conditions, particularly ADHD (19), which resembles sleep deprivation with regard to attentional lapses and a reduced ability to focus (51). Another interesting study found that medication-naive adolescents with ADHD had increased offsets and slopes in their power spectra, which normalized after stimulant treatment (52). In this regard, it is also promising to look at recovery sleep after sleep deprivation, where an increased slow-wave or deep sleep activity was found alongside an increased spectral slope (53). This finding could be interpreted as a shift towards inhibitory activity following wakefulness-induced hyperexcitation.

Importantly, a decrease in the offset and slope of the power spectrum occurs in depression (the opposite of ADHD), and this effect intensifies with an increasing number of depressive episodes experienced throughout one’s lifetime (54,55). Since we demonstrate here that sleep deprivation increases offset and slope, one could conclude that reduced offset and slope could be an additional biomarker for the response to wake therapy besides (prefrontal) theta power, particularly in those patients with multiple episodes (i.e., in chronic/recurrent depressive disorder). These patients with decreased overall neural activity (reduced offset) and diminished slow-wave activity (reduced slope) may be more prone to developing depressive symptoms due to this neurobiological pattern.

### HCTSA

In the HCTSA we found a feature indicating variance to differentiate between sleep deprivation and normal sleep. We discovered a reduced variance or complexity of the signal after sleep deprivation, which might reflect a limited ability of the brain to flexibly react to upcoming stimuli. Previous studies have found a reduced variance (44) or a blunted change in variance in EEG data when responding to stimuli (45). This aligns with our idea of an’overheated engine’, where increased offset and slope reflect uncoordinated neural hyperexcitation. Furthermore, it matches the comparison between ADHD and sleep deprivation, as both sleep deprivation (45) and ADHD (19,52) go along with EEG patterns associated with the inability to process stimuli properly. However, this highly explorative finding needs more research, since research in this area is naturally sparse. Importantly, this brings us to the main take-away of the HCTSA: Despite this highly explorative analysis, no measure performed better than the hypothesis derived theta, highlighting the importance of theta oscillations in sleep deprivation and sleep pressure. Unfortunately, the HCTSA could not be calculated for the mouse data, because the two-hour EEG-recordings proved to be too computationally demanding even on the high-performance computing cluster.

### Complementary analysis of mouse data

The inclusion of mouse data in this study serves as a critical cross-species validation, demonstrating that the electrophysiological and behavioral response to sleep deprivation is a fundamental biological phenomenon rather than a species-specific artifact. This robustness mirrors the human data and reinforces the utility of theta power and aperiodic dynamics as mechanism-based biomarkers. However, a key observation in this comparative approach is a small but characteristic shift in oscillatory frequency between species. Previous studies have shown that functionally equivalent theta is slower in humans compared to rodents (56,57). Interestingly, we also found an increased spectral offset after sleep deprivation, which is in line with the human data. Other studies also discovered a steeper spectral slope after sleep deprivation (48), however, due to small number of subjects it is little surprising that the effect did not reach significance in our analysis. We have to acknowledge the limitations by the sample size and thus future studies should replicate our findings with more subjects. The translational value of these shared markers is underscored by the behavioral and neural parallels between the two species. In both humans and mice, sleep deprivation is a known trigger for mania-like behaviors (26,49). The high degree of similarity in EEG signatures observed here suggests that the increase in theta power and spectral offset after prolonged wakefulness is a conserved mammalian response. Consequently, these markers provide a validated framework for monitoring therapeutic interventions like wake therapy in major depressive disorder, offering a bridge between invasive mechanistic research in mice and clinical application in humans.

## Conclusion

This study systematically investigated the electrophysiological signature of prolonged wakefulness using a robust, multi-methodological (power spectrum, SpecParam, and HCTSA) and cross-species approach across independent EEG datasets. Our findings confirm and extend the understanding of how acute sleep deprivation alters resting-state neural activity, providing critical insights for therapeutic applications. The most salient finding unequivocally establishes theta power as the single most consistent electrophysiological marker of prolonged wakefulness in humans and mice. Theta power was the most reliable feature for differentiating between the control and sleep-deprived conditions across all analyses, with the SVM demonstrating high classification accuracy based on this feature. Even the highly exploratory HCTSA confirmed that no alternative feature outperformed hypothesis-derived theta power as a reliable electrophysiological marker. Beyond traditional oscillatory activity, the analysis of aperiodic components provided novel functional interpretations. The increased power spectral offset suggests a state of global cortical hyperexcitation, while the accompanying steeper spectral slope suggests a relative shift toward slow-wave activity. This simultaneous shift represents an uncoordinated neural hyperexcitation. These consistent and robust markers carry significant translational implications for wake therapy in depression. Based on this mechanism, we propose two potential biomarkers for predicting an individual’s treatment response to wake therapy: (1) Reduced (prefrontal) theta power (a state potentially restored or’reset’ by successful sleep deprivation) and (2) reduced offset and slope of the power spectrum (a state potentially normalized in responders following therapeutic intervention). Future research should focus on validating these proposed EEG markers in clinical cohorts to transition these robust findings into effective clinical tools for personalized chronotherapeutic interventions.

## Materials and Methods

### Participants

The Chinese research group recruited 71 healthy adults (34 females) with an age range of 17 to 23 years (*M* = 20, *SD* = 1.44)(58). In the German study 31 healthy adults (15 females) with a mean age of 24.62 (*SD* = 4.16) years were measured. The nine mice were 12 weeks old when the EEG electrodes were applied during surgery.

**Table 1.** Overview of the analyzed datasets. We used a publicly available dataset from Southwest University, Chongqing, China and the Leibniz Research Centre for Working Environment and Human Factors in Dortmund, which were used as discovery and replication dataset, respectively. Finally, datasets were merged for an even more robust analysis. To validate our approach even across species, we analyzed a dataset from a south Korean group, measured in mice. Regarding the sleep deprivation protocol this corresponds to the human datasets.

| Dataset | Origin | N | Excluded | Species |
| --- | --- | --- | --- | --- |
| Set 1<br>Discovery<br>Dataset | Southwest University,<br>Chongqing,<br>China | 71<br>(34 female) | 6 | Human |
| Set 2<br>Replication<br>Dataset | Leibniz Research Centre<br>for Working Environment<br>and Human Factors,<br>Dortmund, Germany | 31<br>(15 female) | 0 | Human |
| Combined<br>Dataset | Chinese data<br>+<br>German data | 102<br>(final<br>sample:<br>N = 96) | 6 | Human |
| Set 3<br>Dataset for<br>interspecies<br>validation | Korea Institute of Sci-<br>ence and Technology,<br>South Korea | 9<br>(0 female) | 0 | Mouse<br>(C57BL/6 and<br>129S4/SvJae<br>hybrid) |

### Procedure

In both human studies (21,22), participants completed two sessions approximately one week apart: a control sleep session and a 24-hour sleep deprivation session, with the order of sessions randomized. During the sleep deprivation sessions, participants remained continuously awake for 24 hours under the supervision of study personnel. For the sleep control sessions, participants were instructed to have a night of regular sleep. In each case, a fixed time of day was used for testing to minimize the influence of circadian rhythms. Following these sleep conditions, resting-state wake EEG recordings were gathered with a 61-channel EEG system (Brain Products GmbH, Germany) in Chongqing and a 64-channel EEG system (NeurOne, Bittium Corporation, Finland) in Dortmund.

The mouse data were gathered by surgically implanted electrodes with a 32-channel high-density array. Following recovery from surgery, nine days of continuous recording were performed, which included two baseline days, six days of chronic sleep restriction, and one recovery day. To induce sleep deprivation, the mice were placed in cages with a rotating bar mechanism that automatically disturbed their sleep for 20 hours each day. The procedure also featured a specific two-hour high-density EEG session on the first day of restriction to capture high-resolution spatial dynamics of the brain during sleep loss. This was the period we analyzed and compared to the EEG after the baseline nights.

### EEG Preprocessing

Human EEG data from China and Dortmund underwent a standardized preprocessing using an automatic pipeline adapted for resting-state recordings (59) from a previously proposed method(60). This pipeline represents a concatenation of established functions within the Matlab-based EEGLAB toolbox (61), ensuring uniform data quality and high reproducibility across subjects. The preprocessing sequence comprised the following critical steps:

Downsampling the continuous data to 250 Hz, line noise removal to mitigate power line artifacts and robust bad channel rejection. The data was then re-referenced to the average reference across all remaining channels. Artifact removal was primarily achieved through Independent Component Analysis (ICA), with subsequent automated rejection of independent components labeled by a machine learning classifier (ICLabel; (62) as “muscle” or “eye” artifacts. Finally, the cleaned data was subjected to bad segment rejection before being epoched into 2s segments with 1s overlaps, making the data ready for frequency-domain analysis. All preprocessing functions were executed using their default parameter settings. Six of the 102 participants were excluded since they had less than 50 clean trials resulting in a total sample of 96 participants for the analysis. In contrast to the automated human pipeline, the mouse EEG recordings - which featured significantly longer durations of approximately two hours per subject - were cleaned using a semi-automated visual rejection approach based on trial variance. The data was therefore again separated in artificial trials of two seconds. This was conducted using the FieldTrip toolbox to ensure that the higher volume of data per animal was rigorously screened for non-biological artifacts. We systematically identified and removed trials displaying excessive variance across channels. This visual inspection and rejection procedure was applied independently to both the baseline and sleep deprivation conditions. This ensured that only high-quality, artifact-free segments were retained for subsequent frequency-domain analysis and the training of the SVM classifier.

## Analyses

### Statistical analysis

We analyzed two human datasets with eyes-open resting-state EEG, measured by two different groups in China and Germany. In doing so, we were trying to answer different research questions, than the groups collecting the data. Our goal was to identify biomarkers of sleep deprivation to better understand the biological basis of wake therapy and sleeping problems in mental health, primarily depression and non-organic insomnia. To this end, we computed various analyses using different approaches across the combined dataset (Chinese + German data). First, we used the power spectrum to identify oscillations associated with sleep deprivation with a special focus on the theta frequency as a marker for neural plasticity. To identify significant frequency clusters within channels, power spectra were divided into frequency bins of (0.25) Hz width. For each frequency bin, paired-sample t-tests compared the two sleep conditions (first-level statistic). Adjacent frequency bins exceeding a predefined threshold (*p* < 0.05, uncorrected) were grouped into clusters. For each cluster, the cluster-level statistic was computed as the sum of all t-values within that cluster. To assess the significance of each cluster, a Monte Carlo permutation procedure (5000 randomizations) was applied in which the condition labels were randomly exchanged within participants (63). For every permutation, the largest cluster-level statistic was stored, yielding an empirical null distribution. The *p*-value of an observed cluster corresponds to the proportion of permutations in which a cluster statistic of equal or greater magnitude was obtained. On the topographical plots, the color scale represents the original (uncorrected) *t*-values, while the outlined regions indicate sensors belonging to clusters that survived cluster-based permutation correction (cluster-level *p* < 0.05).

Additionally, to this frequentist approach, we also calculated channel-wise Bayesian *t*-tests to compare sleep deprivation with normal sleep. The same spectral bandwidth of 0.25 Hz was used to compute Bayes Factors to investigate the presence and absence of effects. Within these 0.25 Hz windows, the two sleep conditions were compared. We interpret log_10_ Bayes factors between 0 and ½ as not worth mentioning, between 1/2 and 1 as substantial, between 1 and 2 as strong and greater log_10_ Bayes factors as decisive (64). The results of the Bayesian tests are shown in the supplementary material.

To disentangle the homeostatic changes in narrow-band oscillations (periodic activity) from the broadband spectral shifts (aperiodic activity), we utilized the fooof (Fitting Oscillations & One-Over-F) algorithm (also known as SpecParam (47)) integrated into the FieldTrip toolbox (63). This approach addresses the “1/f” nature of the EEG power spectrum, ensuring that changes attributed to specific frequency bands (e.g., theta) are not artifacts of shifts in the underlying broadband slope or offset. Total power spectra were first estimated using a Multi-Taper Fast Fourier Transform (MTMFFT) with a Hanning taper (frequency range: 1–100 Hz, 4s padding). The resulting power spectra were then decomposed into periodic and aperiodic components. The same statistical and machine learning analysis procedure was applied as for the uncorrected spectrum. To characterize the neural dynamics across the entire physiological frequency range the algorithm was fitted from 1 to 100 Hz. We used the “fixed” instead of the “knee” approach to estimate the aperiodic fit. This was done to avoid overfitting, improve interpretability and computational stability. However, it is important to note that slope/exponent estimates depend on the frequency range, aperiodic mode (knee vs. fixed), and referencing.

Both the human and mouse analyses made use of the standard power spectrum and the use of the SpecParam-algorithm to disentangle periodic theta oscillations from aperiodic 1/f dynamics. Both were analyzed using the same settings in the standard and corrected power spectrum. To ensure statistical comparability despite differences in data volume, the mouse analysis incorporated a randomized trial-balancing procedure to equalize trial counts between conditions. Regarding the calculation of the cluster-statistics for the offset and slope in the mouse data only 512 (instead of 5000) permutations were possible due to the low number of subjects (N = 9). Thus, the cluster-statistics were reported based on the maximum possible number of permutations.

### Machine learning Analysis

To assess whether EEG time-series features can discriminate between normal sleep and sleep deprivation, we implemented a supervised classification pipeline. This more complex multivariate machine learning (ML) analysis is using a support vector machine (SVM). First, for each subject the difference of both conditions (sleep deprivation - normal sleep) was calculated, as this analysis would not consider that both conditions were gathered from the same subject (as the paired t-test does). We trained the algorithm in the discovery (Chinese) dataset and re-ran the same analyses in the replication (German) dataset to investigate the stability of the discovered effects. This leave-site-out approach emphasizes the robustness of biomarkers in different recording environments. Finally, to achieve the most robust results possible, we repeated all analyses using the combined dataset. Machine Learning was performed for the standard and corrected power spectrum, slope and offset of the power spectrum and the best performing feature in the HCTSA.

Cross-validation was conducted at the subject level: in each fold, both conditions from a subject were assigned jointly to the training or test set. We used 10-fold cross-validation repeated 30 times with different random partitions. Importantly, all data from a single subject is kept together in either the training or test set, so that we ensure that the machine learning model is learning to identify biomarkers of sleep deprivation rather than just identifying the unique brain signature of a specific person. For each feature, we recorded two performance metrics: overall classification accuracy and the area under the receiver operating characteristic (ROC) curve (AUC). Again, the machine learning pipeline was also applied to both the uncorrected and corrected power spectrum, just as the statistical analyses.

The machine learning approach in the mouse was conducted with the same settings for the classifier. However, the main difference lies in the architecture of the training and test sets. The human machine learning approach focused on inter-subject classification, training a model on a large discovery cohort and validating it on an independent replication cohort to ensure the features generalized across different people. In contrast, the mouse analysis utilized with-in-subject, single-trial classification. Because the mouse dataset provided continuous long-term recordings (approx. two hours), we could leverage hundreds of trials per individual to train high-precision models tailored to the specific neurophysiology of each mouse before aggregating the performance results. For every subject, we identified the condition with fewer clean trials and randomly sub-sampled the other condition to match it perfectly. This ensured that the mouse models were trained on equally sized training sets. As in the human data, the machine learning pipeline was applied to the standard and corrected power spectrum.

### HCTSA

For the HCTSA analysis (29) 7288 features were analyzed and the feature that differentiated best between the two sleep conditions was extracted. The analysis was executed using the high-performance computing cluster, PALMA II, which is situated at the University of Münster. All channels were tested with a univariate paired *t*-test (with cluster-based multiple comparisons correction) for differences in this feature between the sleep control condition and sleep deprivation. As in the previous analyses we also used a SVM to analyze the extracted feature. The feature that differentiated best between sleep deprivation and normal sleep was feature 1592 named “CO_Embed2_Basic_tau_incircle_2”. This represents the fraction of points in a 2D time-delay embedding plot that fall inside a circle centered at the origin with a squared radius of 2. A low value of this feature means that in the decorrelation timescale of the series, the joint distribution is spread far from the origin. This suggests higher variance, heavy-tailed behavior, or richer nonlinear structure in the dynamics, rather than a compact, noise-like cloud near 0,0. In simpler terms, it’s a measure of how densely the time series data is clustered around the central point (0,0) in this special plot. It can also be seen as a measure of the variability or complexity in EEG data after sleep deprivation compared to normal sleep.

The HCTSA could not be calculated for the mouse data due to the long duration (two hours) of the EEG data. Even on the high-performance computing cluster, the computation time was so extensive that the task could not be completed.

### Control Analyses of Mouse Data

Since sleep deprivation can induce “mania-like behavior” not only in humans but also in mice a control analysis of the mouse data is necessary because the mice were allowed to move during the EEG recording. Thus, it could be that the EEG changes (in particular theta power) could be trivially explained by increased locomotor activity or movement-related artifacts, rather than reflecting genuine neural state changes. To solve this problem, we computed control analyses by using the likewise measured accelerometer data. First, we compared the movement data after the night of normal sleep with the sleep deprivation condition. This revealed no significant effect of sleep deprivation on overall movement (*t*(8) =-1.93, *p* = 0.09), however, the Bayes Factor (*BF_10_*= 5.42) tends to favor H_1_, i.e., more movement after sleep deprivation.

For this reason, we conducted a second control analysis, correlating the accelerometer data with theta power, our most reliable predictive feature, to demonstrate that the observed effects are due to genuine changes in brain activity rather than altered locomotion. This showed no association between the accelerometer data and theta power (*r*(8) = 0.057, *p* = 0.29). Importantly, the Bayes Factor also strongly favors H_0_ (*BF_01_* = 0.22). Accordingly, it is unlikely that movement is a confounding factor in the calculated analyses. Respective figures are shown in the supplementary material. For the human data this analysis is not necessary as the subjects were not allowed to move during the resting-state EEG.

## Data and code availability

The Chinese human EEG data is available at: 10.18112/openneuro.ds004902.v1.0.8 and the German human EEG data was made available upon request. The mouse EEG data from Korea is available at: 10.12751/g-node.mvi0dp. The analysis pipeline used for this paper is stored at: doi: 10.12751/g-node.5tzhqw

## Acknowledgements

This work was funded by the consortium grant Trajectories of Affective Disorders from the German Research Foundation (DFG) SFB/TRR 393 (project grant no 521379614).

UD was funded by the German Research Foundation (DFG, grant FOR2107, DA1151/5-1, DA1151/5-2, DA1151/9-1, DA1151/10-1, DA1151/11-1 to UD; SFB/TRR 393, project grant no 521379614) and the Interdisciplinary Center for Clinical Research (IZKF) of the medical faculty of Münster (grant Dan3/016/26 to UD).

The work of JA was supported by NEURON-ERA-NET 2021 (MINERVA, DLR FKZ 01EW2209B), the German Research Foundation (DFG) EXC 1003 (Grant FF-2014-01 Cells in Motion–Cluster of Excellence, Münster, Germany), the DFG FOR2107 AL1145/5-2, IMF DL121204 and in part by the SFB/TRR 393-1 consortium from the DFG (subproject A05).

## Author Contributions

T.K. developed the methodology, analyzed the data, wrote the initial draft of the manuscript and created the figures; L.P. reviewed and edited the manuscript; O.A. reviewed and edited the manuscript; S.G. reviewed and edited the manuscript; C.K. reviewed and edited the manuscript; I.W. reviewed and edited the manuscript; M.A.S. acquired parts of the data and reviewed and edited the manuscript; T.R. reviewed and edited the manuscript; J.A. acquired funding and reviewed and edited the manuscript; U.D. acquired funding and reviewed and edited the manuscript; P.R. acquired funding and reviewed and edited the manuscript; J.G. developed the methodology, acquired funding and reviewed and edited the manuscript.

