## Supplementary Materials for "Multi-Methodological Characterization of Sleep Deprivation: From Standard EEG Power Spectra to Aperiodic Dynamics in Humans and Mice"

### **1.1. Bayesian statistics**

**Standard power spectrum**

As in the frequentist approach, Bayesian *t*-tests revealed significant increases in theta power following sleep deprivation (Figure SM1A), with strongest effects over occipital and central electrodes (Figure SM1B). The maximal log_10_BF of 2.85 was reached at 5.5 Hz, which can be classified as decisive evidence^1^.

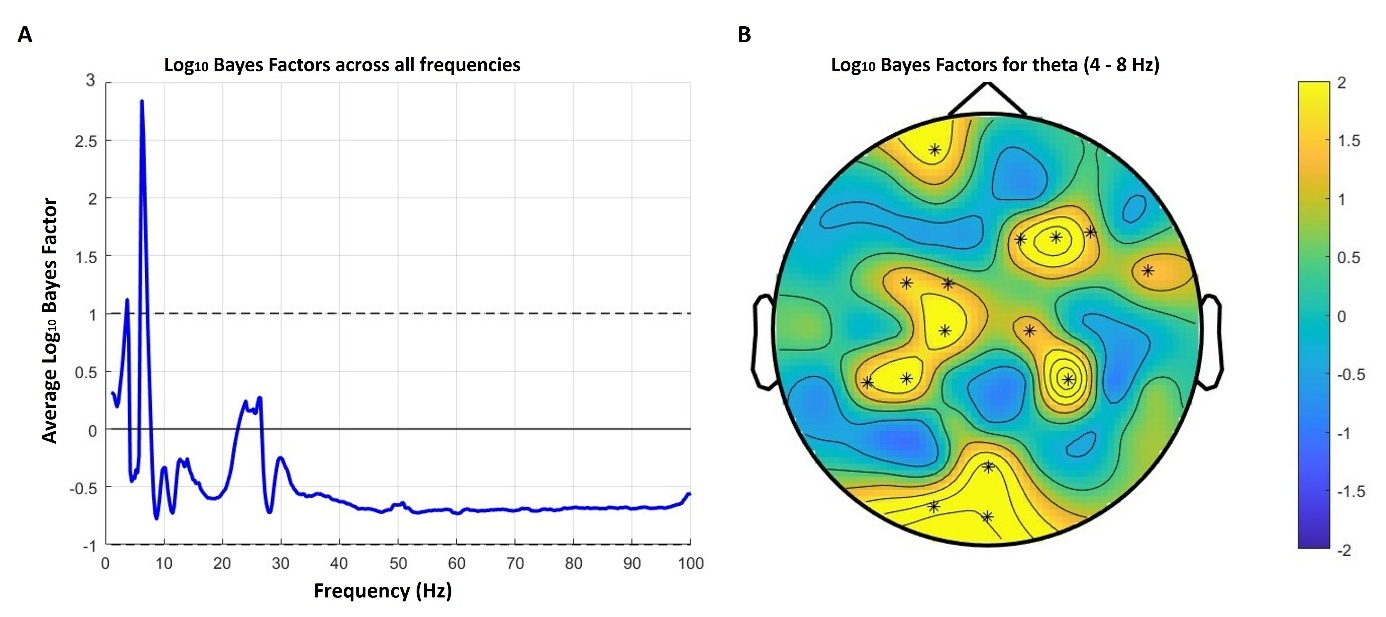
*Figure SM1.* **A**. Logarithmized Bayes Factor averaged over all electrodes in each frequency differentiating between normal sleep and sleep deprivation. **B**. Topography plot indicating Bayes Factors testing normal sleep against sleep deprivation in the theta frequency range.

**Corrected power spectrum**

The results in the corrected power spectrum are similar to the ones in the normal power spectrum with greatest effects in the theta range (maximal log_10_BF = 2.85 at 8 Hz; Figure SM2A). The Bayesian approach also showed an effect in the gamma range (peaking at 52.25 Hz, log_10_BF = 1.42; strong evidence^1^), indicating higher gamma power after normal sleep compared to sleep deprivation.

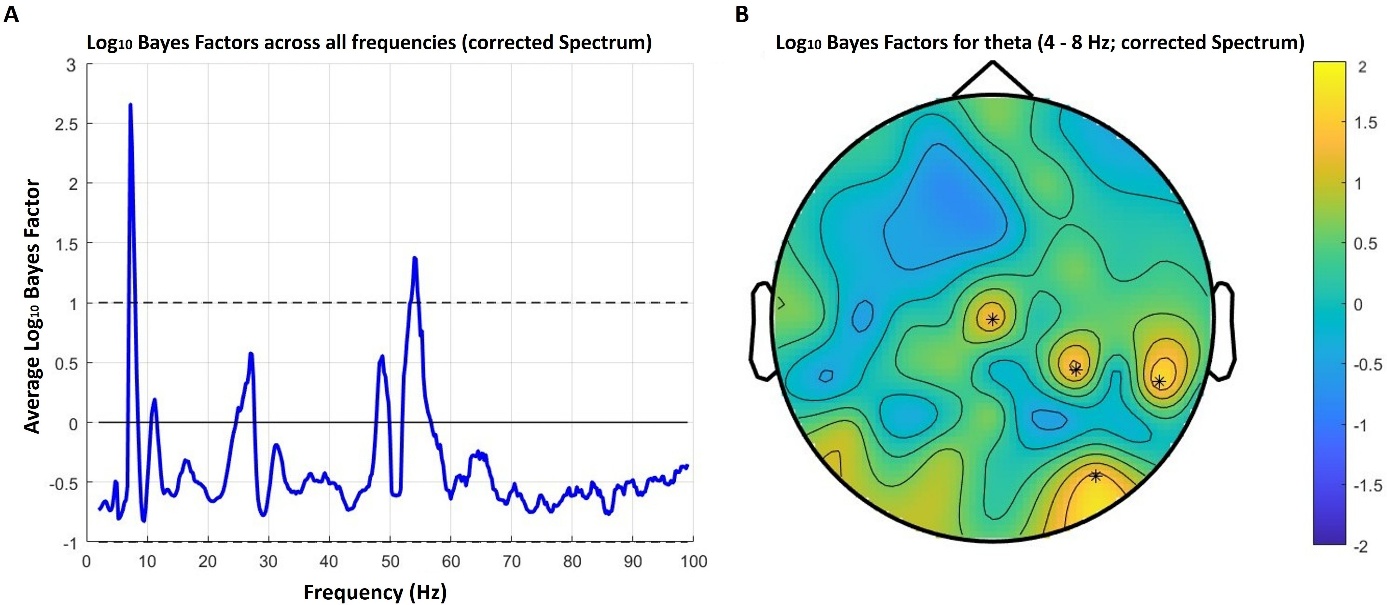

*Figure SM2.* **A.** Logarithmized Bayes Factor averaged over all electrodes in each frequency differentiating between normal sleep and sleep deprivation in the corrected power spectrum. **B.** Topography plot indicating Bayes Factors testing normal sleep against sleep deprivation in the theta frequency range (corrected Spectrum).

**Offset and Slope**

Comparing the offset and slope of the power spectrum revealed significant differences, particularly with regard to the offset, which was evident across a range of electrodes distributed across the entire scalp. Nevertheless, log_10_ Bayes Factors indicated that the strongest effects of sleep deprivation on slope were over the posterior electrodes.

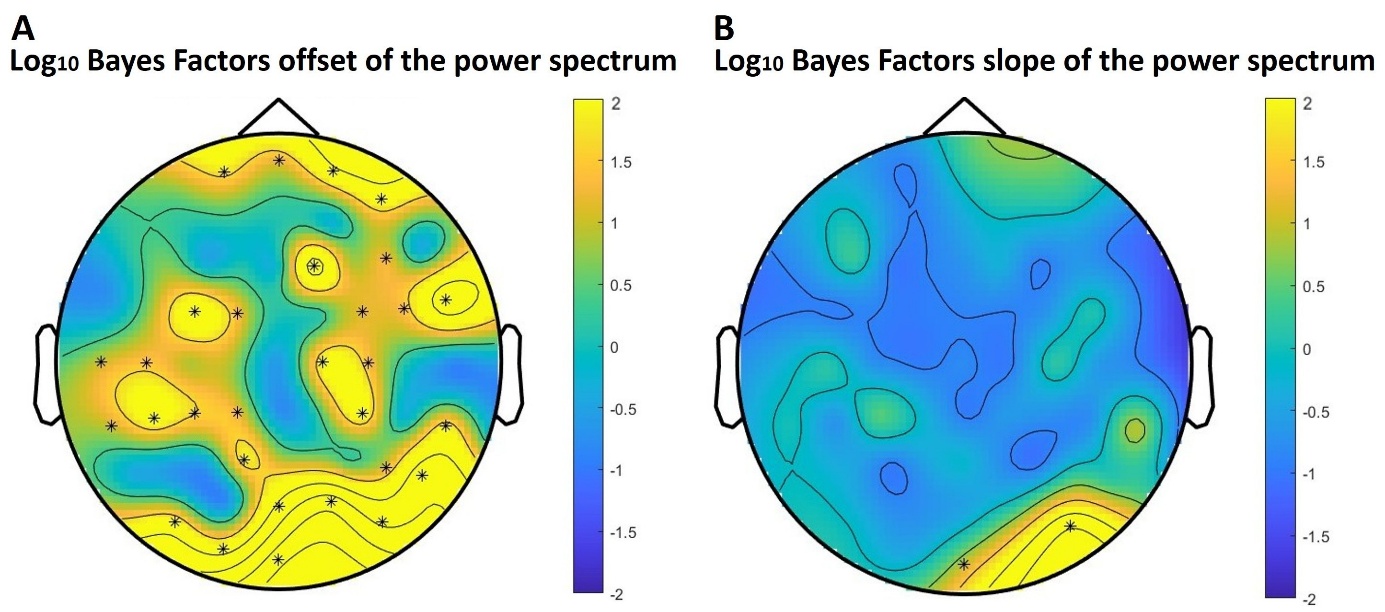

*Figure SM3.* **A.** Topography plot indicating log_10_ Bayes Factors testing offset of the fooof-corrected power spectrum after normal sleep versus sleep deprivation. **B.** Topography plot indicating log_10_ Bayes Factors testing slope of the corrected power spectrum after normal sleep versus sleep deprivation.

**HCTSA**

HCTSA feature 1592  “CO_Embed2_Basic_tau_incircle_2”, was best at discriminating between wake EEG after normal sleep and sleep deprivation. This can be understood as measure of variance or complexity of the signal. The topographies indicate a reduced variance after sleep deprivation compared to normal sleep over various electrodes.

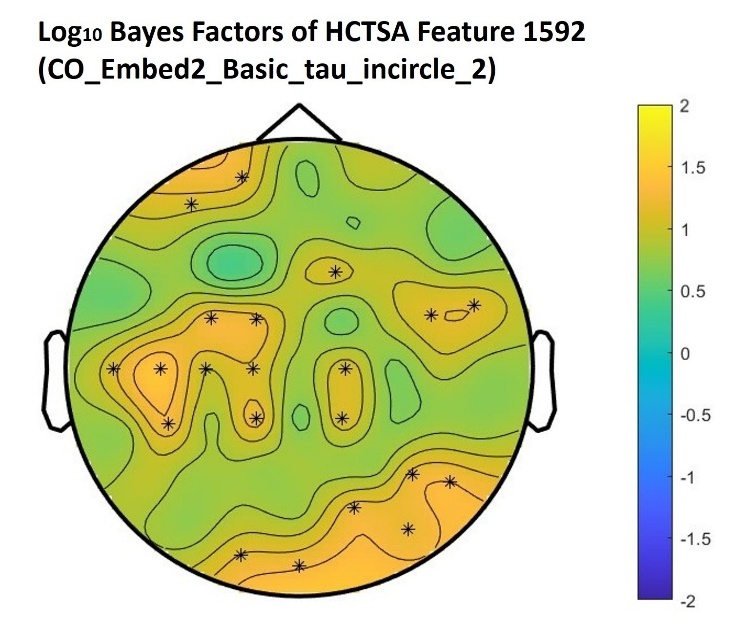

*Figure SM4.* Topography plot indicating log_10_ Bayes factors were used to test the HCTSA feature 1592 after normal sleep versus sleep deprivation.

### **1.2 Control Analysis for Movement Artifacts in Mice**

To ensure that brain activity was not confounded with locomotion we computed control analyses to make sure that theta power was not confounded with movement. It is known that theta power in rats can be modulated by movement; however, this is primarily associated with hippocampal rather than cortical theta rhythms^2–4^.

Therefore, we compared the accelerometer data, which was recorded in parallel to the analyzed EEG sessions, and tested whether the wake EEG in the normal sleep condition differs from the sleep deprivation condition. Although the frequentist statistics did not show a significant difference, (*t*(8) = -1.93, *p* = 0.09), the Bayesian statistics rather supports (*BF_10_* = 5.42) more movement in the sleep deprivation condition (Figure SM5). This unclear pattern demands further control analyses ensuring that EEG (especially theta power) is not confounded by movement artifacts.

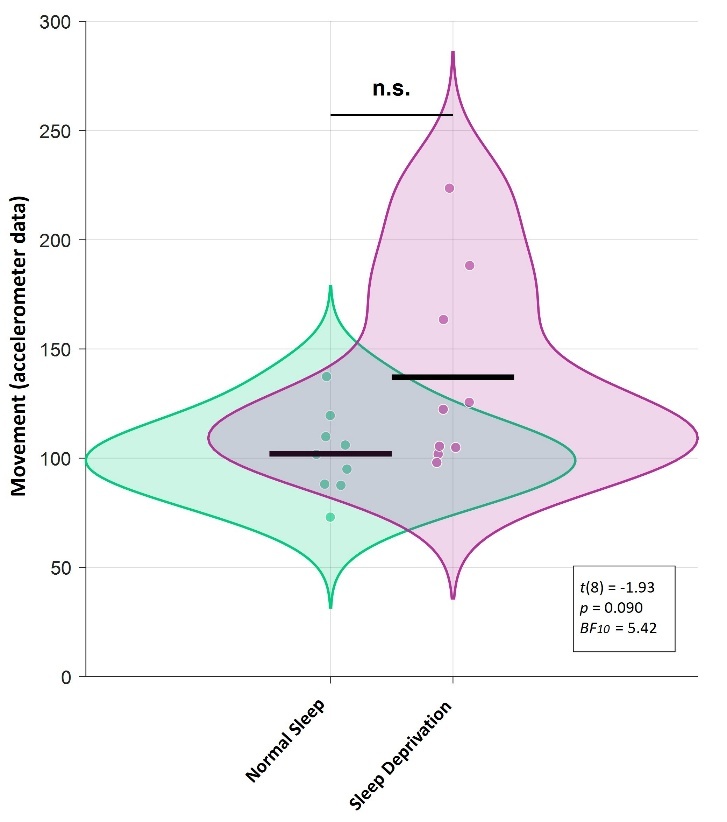

*Figure SM5.* Accelerometer data after a night of normal sleep versus sleep deprivation in mice. These were recorded in parallel to the analyzed EEG data.

To make sure that our statistical and machine learning effects are due to genuine changes in brain activity and not induced by locomotion, we tested the association between theta power and the accelerometer data. This showed no significant association in the frequentist approach (*r*(8) = 0.057, *p* = 0.29) and the Bayes Factor (*BF_01_* = 0.22) clearly supports H_0_, i.e., no association between movement and theta power. Thus, we can rule out movement as a confound in our analyses.

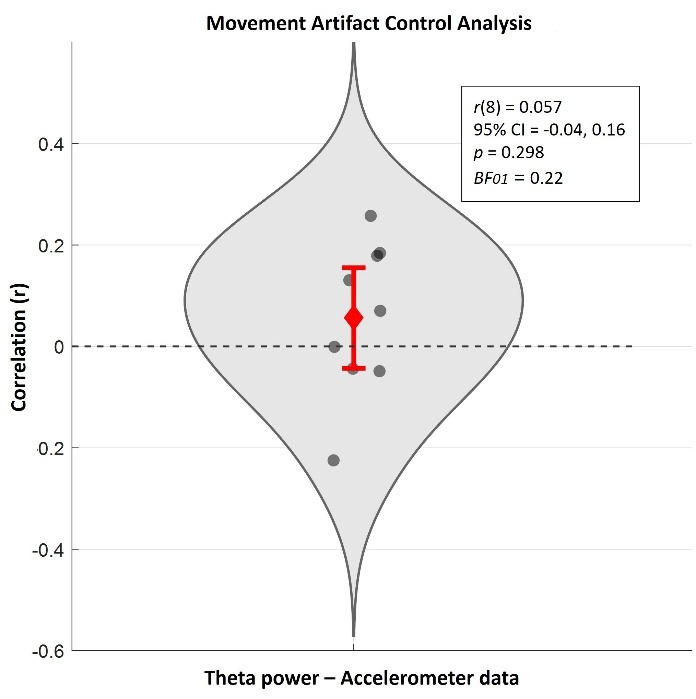

*Figure SM6.* Correlation of theta power and movement in mice wake EEG.
